# From Brain Microstructure to Dynamics: Linking Grey and White Matter Architecture to Propagation Delays

**DOI:** 10.64898/2026.06.12.731878

**Authors:** Svetla Manolova, Carolyn McNabb, Eirini Messaritaki, Eleonora Lupi, Alessandro Daducci, Marco Palombo, Krish D. Singh, Derek K Jones, Mara Cercignani, Matteo Mancini

**Affiliations:** Cardiff University Brain Research Imaging Centre (CUBRIC), School of Psychology, Cardiff University, Cardiff, Wales, United Kingdom; Department of Brain and Behavioral Sciences, University of Pavia, Pavia, Italy; Diffusion Imaging and Connectivity Estimation (DICE) Lab, Department of Computer Science, University of Verona, Verona, Italy; School of Computer Science and Informatics, Cardiff University, Cardiff, United Kingdom; Centro Ricerche Enrico Fermi, Rome, Italy

## Abstract

Understanding the relationship between neural dynamics and underlying brain structure, and in particular the impact of the latter on conduction delays, remains a core question in neuroscience. In humans, we primarily access these phenomena at the macroscale through neurophysiological measures of propagation. At this scale, signal propagation is reflected in measurable delays arising from both white matter (WM) axonal conduction and grey matter (GM) synaptic and integrative processes.

While specific WM features have been explicitly linked to single-axon conduction delays, no clear models link GM microstructural properties to large-scale propagation. To this aim, we combined advanced MRI microstructural modelling with resting-state MEG to link brain structure to signal propagation in a sample of 94 healthy controls. WM metrics included axonal diameter, myelination (g-ratio, myelin water fraction), and tract length, while GM was characterised using the SANDI model and cortical thickness. Propagation delays were quantified using the neuronal avalanches framework.

Our findings indicated significant associations between all of WM and GM metrics and the propagation delays. When attempting to use our W/GM metrics we were able to predict up to ~26% of the observed delay using linear regression modelling. We also found associations between frequency-specific propagation dynamics and tract length. Finally, using a canonical correlation analysis we demonstrated multivariate coupling between our W/GM metrics and the MEG frequency-specific propagation delays. Our findings provide evidence for the link between tissue structure and large-scale neural dynamics, supporting the development of biologically grounded models of signal propagation.

## Introduction

Understanding how brain structure influences brain function remains a fundamental question in neuroscience (Mancini et al., 2021). Long-range coordination and synchronisation between distributed grey matter (GM) regions is supported by white matter (WM) pathways and timely neural signal transmission, which introduces time delays (Bargmann & Marder, 2013; Seguin et al., 2023). At the macroscopic level we observe *propagation delays* - the time associated with the transmission of activity between structurally connected GM regions, which not only reflects axonal conduction (and associated conduction delays) but also synaptic transmission and post-synaptic integration. Within WM, key determinants of signal transmission such as axonal length, diameter and myelin sheath thickness have been well described (Rushton, 1951; Drakesmith et al., 2019). Nevertheless, there are currently no explicit models linking GM microstructural properties to macroscale propagation dynamics. Although cable theory and related biophysical models - which treat dendrites as electrical cables - provide principled predictions regarding how neuronal morphology shapes signal integration and transmission (Rall, 1962; Hodgkin & Huxley, 1952), it remains unclear how microstructural features like neurite density and cell body organisation together with WM cellular structure jointly shape macroscale propagation dynamics.

Diffusion MRI (dMRI) provides a non-invasive means of probing tissue microstructure by characterising the random motion of water molecules within biological tissue, which is constrained by cellular features such as membranes, neurites, and cell bodies. As a result, dMRI is sensitive to the underlying microstructural architecture of both GM and WM. Advances in biophysical diffusion modelling have enabled increasingly specific estimates of tissue properties, with models such as neurite orientation dispersion and density imaging (NODDI) providing metrics related to neurite density and orientation dispersion (Zhang et al., 2012), and more recent frameworks such as soma and neurite density imaging (SANDI) extending this approach to estimate GM-specific features including cell body signal fraction, apparent soma radius, and neurite signal fraction (Palombo et al., 2020). In parallel, dMRI also enables tractography-based reconstruction of WM pathways, providing macroscale estimates of structural connectivity (Dell’Acqua et al., 2024; Hagmann et al., 2010). Complementary MRI techniques can further characterise tissue composition, including myelin-sensitive methods such as multicomponent relaxometry (MacKay & Laule, 2016; Mancini et al., 2020). Together, these approaches provide in vivo proxies of the biological architecture that may support large-scale brain dynamics.

In this work we aimed to leverage a multimodal framework that combines advanced microstructural imaging with resting-state magnetoencephalography (MEG) to study how brain microstructure constrains brain function. To quantify propagation delays we implemented the neuronal avalanches framework (Shriki et al., 2013). Neuronal avalanches are a data-driven method of characterising propagation dynamics. Spontaneous brain activity is represented by spatiotemporal patterns, observable as large over threshold deflections in electrophysiological signals. Importantly, this framework allows us to estimate delays by tracking the latency of signal deflections between structurally connected GM regions during each avalanche. Using this approach, we can construct a delay matrix that reflects whole-brain propagation timings at the millisecond scale, making this suitable for quantifying inter-regional transmission timings. Previous studies using the avalanches framework have relied on coarse tractography-derived measures, such as streamline count, which lacks biological specificity (Sorrentino et al., 2021).

Here we address this gap by leveraging advanced microstructural WM and GM modelling and MEG-derived propagation delays using the avalanches framework. We derived WM metrics including tract length using microstructure-informed tractography, axonal radius, myelin water fraction and g-ratio. We also estimated GM structural metrics such as soma (incl. radius) and neurites signal fractions using SANDI. Finally, we quantified whole-brain propagation delays using resting state MEG, restricting the analysis only to structurally connected regions. This multimodal approach has the potential to provide a new framework for exploring how different structural metrics are associated with macroscopic signal propagation.

Through integrating advanced MRI microstructural modelling with MEG-derived propagation dynamics, this work provides evidence that microstructural organisation contributes to large-scale dynamics. These findings have the potential to establish a foundation for future studies investigating how microstructural architecture shapes macroscale neural communication in health and disease.

## Methods

### Participants

Ninety-four healthy participants (mean age [SD]: 22.91 [3.44]; female/male ratio = 56/38) were recruited as part of The Welsh Advanced Neuroimaging Database (WAND), an integrated large-scale, multi-modal imaging database comprising MRI, MEG and behavioural data from healthy volunteers (McNabb et al., 2025).

Participants were safety screened for both MRI and MEG to confirm study eligibility in accordance with Cardiff University’s MRI and MEG safety guidelines. All participants included in this analysis underwent both the MRI and MEG scanning sessions.

Written informed consent was obtained from each participant in accordance with the ethical approval for this study, granted by Cardiff University’s School of Psychology ethics committee.

### MRI Acquisition and Preprocessing

MRI data were acquired on a 3T Connectom scanner (Siemens Healthineers, Erlangen, Germany), equipped with ultra-strong 300mT/m magnetic field gradients, enabling high b-value diffusion imaging and enhanced microstructural sensitivity. The MRI imaging session included:

- **T1-weighted structural MRI (MPRAGE);**
- **Multi-component relaxometry (McDESPOT)** for deriving myelin water fraction;
- **Multi-shell diffusion MRI data with a fixed diffusion time** (Δ=24ms) for generating tractograms and fitting microstructural models;
- **Multi-shell diffusion MRI data with variable diffusion time (AxCaliber protocol)** for estimating axonal radii.

All the data acquisition protocols and processing steps are fully documented in the WAND data descriptor; therefore, we only summarise the key steps here. Readers are referred to our previous work (McNabb et al., 2025) for the full acquisition parameters, gradient schemes, quality-control procedures and data preprocessing pipeline steps.

### Structural MRI

The anatomical data were acquired using an MPRAGE sequence at 1mm isotropic resolution. Data were processed and segmented using the Desikan-Killiany atlas from Freesurfer 5.3.0 (Dale et al., 1999; Desikan et al., 2006) - from the atlas, 82 regions of interest (ROIs) were extracted (68 cortical, 14 subcortical). Cortical thickness (CT) for each ROI was also estimated.

### Multi-component relaxometry

McDESPOT data were preprocessed using the standard three component relaxometry fit using QUantitative Imaging Tools (QUIT) and aligned to obtain myelin water fraction (MWF) maps (Cabeen et al., 2018; Wood, 2018). The MWF maps were aligned to the T1-weighted volume first using FSL’s flirt with the correlation ratio as a cost function (Jenkinson & Smith, 2001; Jenkinson et al., 2002). The affine transformations from the native space to the diffusion space and from the McDESPOT space to native space were concatenated and were then used to align the MWF maps to the diffusion data, again using FSL’s flirt.

### Diffusion data

All diffusion-weighted data preprocessing included denoising and corrections for signal drift, susceptibility, eddy current-induced distortions, gradient non-uniformity and Gibbs ringing following the preprocessing pipeline described in our previous work (McNabb et al., 2025; Cordero-Grande et al., 2019; Tournier et al., 2019; Veraart, et al., 2016a; Veraart et al., 2016b; Andersson et al., 2003; Smith et al., 2004; Kellner et al., 2016).

### Tractography

Using boundary-based registration, we estimated an affine transformation from the diffusion data to the FreeSurfer anatomical space and then we aligned the anatomical data using the inverse of this transformation (Greve & Fischl, 2009; Jenkinson & Smith, 2001; Jenkinson et al., 2002). For the diffusion data with a fixed diffusion time we used multi-shell multi-tissue spherical deconvolution as implemented in MRtrix3 to estimate the fibre orientation distribution (Jeurissen et al., 2014; Tournier et al., 2019) and then anatomically constrained tractography to probabilistically reconstruct 5 million WM matter pathways, cropping each streamline at the GM/WM interface (Smith et al., 2012). We then used the Convex Optimization Modelling for Microstructure Informed Tractography (COMMIT) framework, which combines tractography with microstructural features of the underlying reconstructed tissue with the aim of improving the anatomical accuracy of the reconstructed tracts (Daducci et al., 2014). We used the stick-zeppelin-ball model from COMMIT to generate estimates of the contribution of each streamline to the global fit of the model, which we then used as a means of pruning each tractogram – streamlines that had an effective contribution of 0 to the global fit were deemed improbable and filtered out. The pruned tractogram was used in all the subsequent analyses.

### Axonal Radius

The diffusion data with the variable diffusion time (AxCaliber protocol) were fitted using the COMMIT-AxSize framework which estimates bundle-specific axonal radii (Barakovic et al. 2021). We fitted the Cylinder-Zeppelin-Ball model to estimate the axonal radius weights for each streamline’s contribution to the global model fit using the AxCaliber data with diffusion time of 42ms only. The calculated weights were then used to generate a weighted mean radius for each WM bundle, which was then used to make an axonal radius connectome for each participant.

### G-ratio

To obtain the average g-ratio per bundle for each participant we firstly ran AMICO to fit the neurite orientation dispersion and distribution imaging (NODDI) model to the multi-shell diffusion data with the fixed diffusion time from which we obtained the neurite density (NDI) and isotropic fraction (ISO) using all the data shells (Daducci et al., 2015; Zhang et al., 2012). From this data we also estimated the diffusion tensor from which we derived the fractional anisotropy (FA) map using volumes with b=0 s/mm^2^ and b=1200 s/mm^2^. The FA map was used to mask out voxels in which FA<0.8 to ensure only voxels with high anisotropy were included. The myelin volume fraction (MVF) can be assumed to be linearly proportional to MWF via a constant alpha. Following a previously proposed approach (Cercignani et al., 2017), we derived alpha by setting the g-ratio equal to 0.7 in the splenium of the corpus callosum which we segmented using TractSeg (Wasserthal et al., 2018). After obtaining the MVF map we then calculated the axonal volume fraction (AVF) and consequently the g-ratio using established formulas and code (Stikov et al., 2015; Lu et al., 2025). For each subject we then generated a connectivity matrix with the average g-ratio per bundle.

### Grey Matter Microstructure

We also fitted the soma and neurite density imaging (SANDI) model to the multi-shell diffusion data with a fixed diffusion time (using all shells) to obtain the neurite fraction (NF), soma fraction (SF) and soma radius (SR) (Palombo et al., 2020). The SANDI maps were fitted using the SANDI toolbox (https://github.com/palombom/SANDI-Matlab-Toolbox-Latest-Release). The SANDI maps were linearly aligned to the Freesurfer anatomical space and the average NF, SF and SR for all the ROIs were extracted. For a representative example of the generated MWF, tract length, axonal radius, g-ratio and SANDI maps for one participant, see Figure 1.

**Figure 1.**
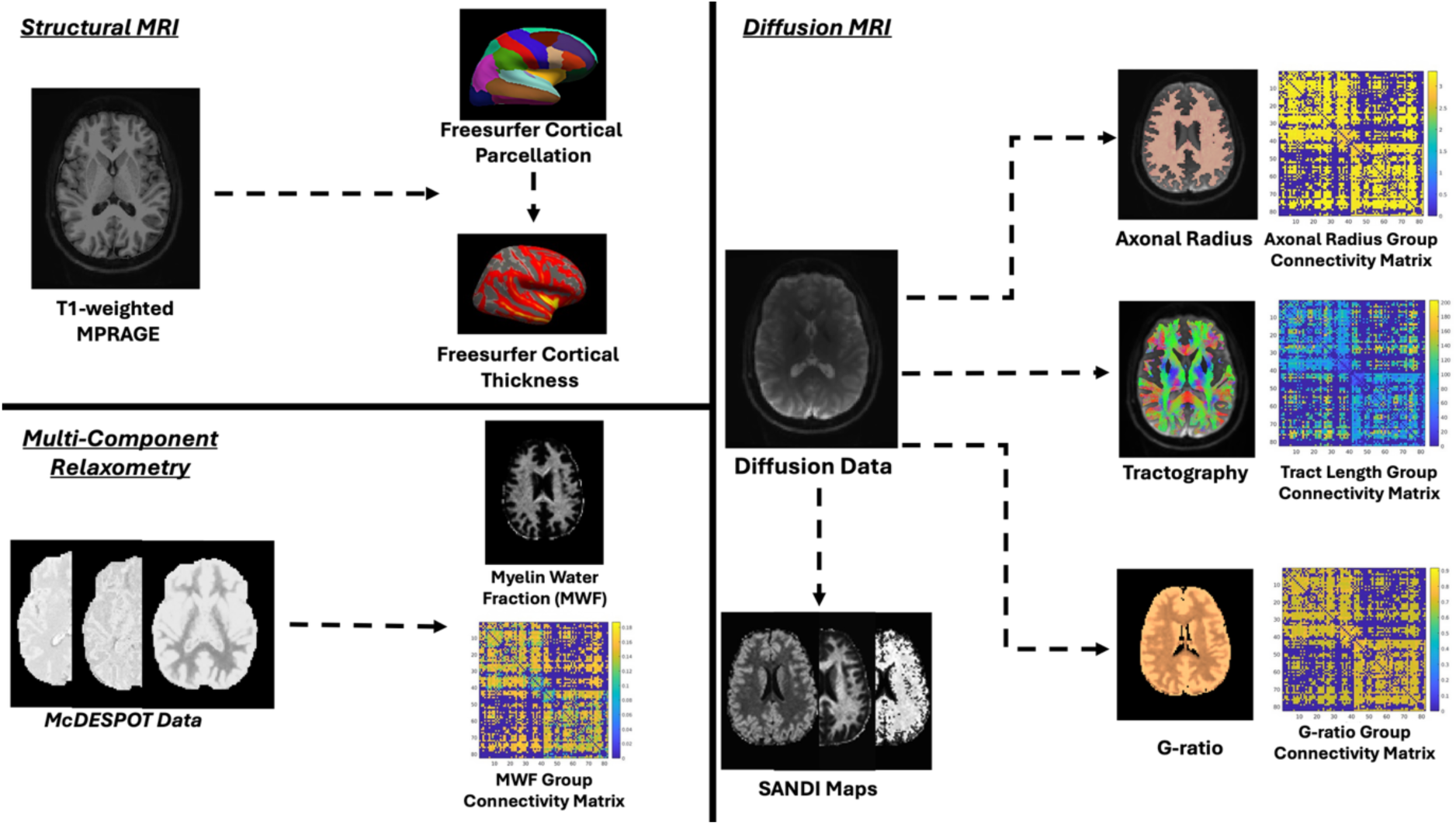
Visual representation of the generated cortical parcellation and structural maps for one representative participant. The “Structural MRI” panel demonstrates the Freesurfer cortical parcellation and the derived cortical thickness map. The “Multi-Component Relaxometry” panel demonstrates the preprocessed McDESPOT data and the derived myelin water fraction (MWF) maps, including the group connectivity MWF matrix. To estimate the MWF index for each bundle we took the mean MWF along each streamline, after which we took the average MWF of all streamlines forming a bundle. Finally, the “Diffusion MRI” panel shows the microstructural maps derived from the diffusion data, including the group connectivity matrices.

### Connectivity Matrices

For each participant we constructed microstructure-weighted connectivity matrices using the parcellated GM regions as nodes and our WM and GM metrics as edge-weights. From our WM metrics we derived the following matrices for each participant: 1) tract length; 2) bundle-average MWF; 3) bundle-specific axonal radius and 4) bundle-average g-ratio. For our GM metrics we used the sum between the GM metric values of each pair of connected ROIs, thus deriving the following matrices: 1) cumulative cortical thickness; 2) cumulative neurite fraction; 3) cumulative soma fraction and 4) cumulative soma radius. As additional GM microstructure-weighted matrices, we also computed the apparent number of cells and the total surface volume (Carriero et al., 2026), as detailed in the supplementary materials. All matrices were thresholded at more than 4 streamlines per connection. Group-consensus matrices were then computed using a threshold of 60% streamline connection prevalence as a trade-off between false positives and false negatives (de Reus et al., 2013).

### MEG Acquisition and Data Processing

MEG acquisition and preprocessing also followed the procedures described in the WAND data description (McNabb et al., 2025), therefore only the key steps relevant to the analyses in this paper are summarised.

### MEG Data Acquisition

Whole-head MEG resting-state data were acquired using a 275-channel CTF radial gradiometer system (CTF Systems Inc., MEGIN, Helsinki, Finland) with a sampling rate of 1200 Hz. Twenty-nine reference channels were recorded to aid with noise cancellation (Vrba & Robinson, 2001). Using a Polhemus device, head digitisation was performed prior to participants entering the magnetically shielded room.

Participants were seated upright in front of a screen with their head supported by a chin rest to minimise movement artefacts. They were asked to keep their eyes open and fixated on a white dot on a grey background. The resting-state acquisition was 10 minutes. Further details on the acquisition process and hardware can be found in the WAND paper.

### MEG Preprocessing and Source Reconstruction

All the datasets were band-pass filtered between 1-150Hz and epoched into 2s-windows. The data were then visually inspected using DataEditor (CTF Systems Inc., MEGIN, Helsinki, Finland) and 2s-epochs containing major artefacts (e.g., eye-blinks, muscle motion etc.) were excluded from further analysis.

Epoched data were downsampled to 512Hz, then source localized using FieldTrip version 20161011 (Oostenveld et al., 2011), with an LCMV beamformer. First the individual’s skull-stripped MRI was spatially normalised to a 2-mm isotropic skull-stripped MNI template brain. This allowed the warping of a 2-mm isotropic source-space for beamforming, including atlas definitions, back to the original individual’s MRI. A head-model in this space was constructed using a single-shell forward model (Nolte, 2003). For each dataset, a global covariance matrix was constructed in a broadband frequency range (1-48 Hz), as well as delta (1-4 Hz); theta (4-8 Hz); alpha (8-13 Hz); beta (13-30 Hz) and gamma (30-48 Hz). Within the source-space, beamforming was performed using a single representative voxel (centroid) in each of the 82 ROIs of the Desikan-Killiany atlas (Desikan et al., 2006). At each of these locations, epochs were concatenated to generate a continuous virtual-sensor time course for each representative voxel by multiplying unit-norm weights for these 82 locations by the data, bandpassed into the required frequency range.

### Neuronal Avalanches

Neuronal avalanches were estimated by computing the z-score of the activity timeseries for each of the 82 MEG virtual sensors and identifying events exceeding a threshold of 3 standard deviations (Shriki et al., 2013). An avalanche is assumed to start whenever one or more timeseries go over threshold and concludes when no timeseries are over threshold. Time was discretized into bins of width Δt, defining consecutive time steps. To determine the optimal bin size for each frequency band, we calculated the branching parameter (σ) (Shriki et al., 2013). The neuronal avalanches are described by a critical branching parameter which is used to identify the critical point from which activity starts propagating in a cascade. The branching parameter represents the average ratio between the number of activations in consecutive time steps. To ensure that the system is in a critical state σ needs to equal 1 as this gives rise to the propagation phenomenon (Shriki et al., 2013).

The branching parameter was derived through calculating the ratio between the number of events in the current time bin to the number of events in the subsequent bin. This was done for the total number of identified avalanches for each subject. The ratio was then geometrically averaged across all the avalanches for each subject. The resulting formulas were (Sorrentino et al., 2022):

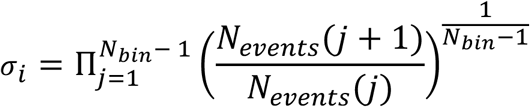

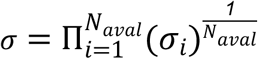

σ*_i_* represented the branching parameter of the current i_th_ avalanche, N_bin_ was the total number of bins in the avalanche, N_events_(j) was the overall number of active events in bin (j), N_aval_ represents the total number of avalanches.

To ensure that avalanche dynamics were evaluated near the critical regime, we estimated the branching parameter for each MEG frequency band. We first randomly selected 70 participants and, for each band, evaluated five candidate bin sizes (1–5 samples), choosing the value that produced a branching parameter closest to 1. Although all tested bin sizes yielded values near criticality, we selected the bin size for each frequency band with the smallest absolute deviation from 1 and applied this value to the full dataset. This procedure resulted in the following optimal bin sizes: whole-signal: 3 samples; delta: 5 samples; theta: 2 samples; alpha: 2 samples; beta: 4 samples; gamma: 5 samples.

Delays were computed within an avalanche counting the number of timesteps between the beginning of the avalanche in a given region and subsequent over-threshold activity of any other region and multiplying by the MEG time resolution (see Figure 2.). A delay connectivity matrix was then built for each subject, averaging over the avalanches, and compared with the WM and the GM structural matrices.

**Figure 2.**
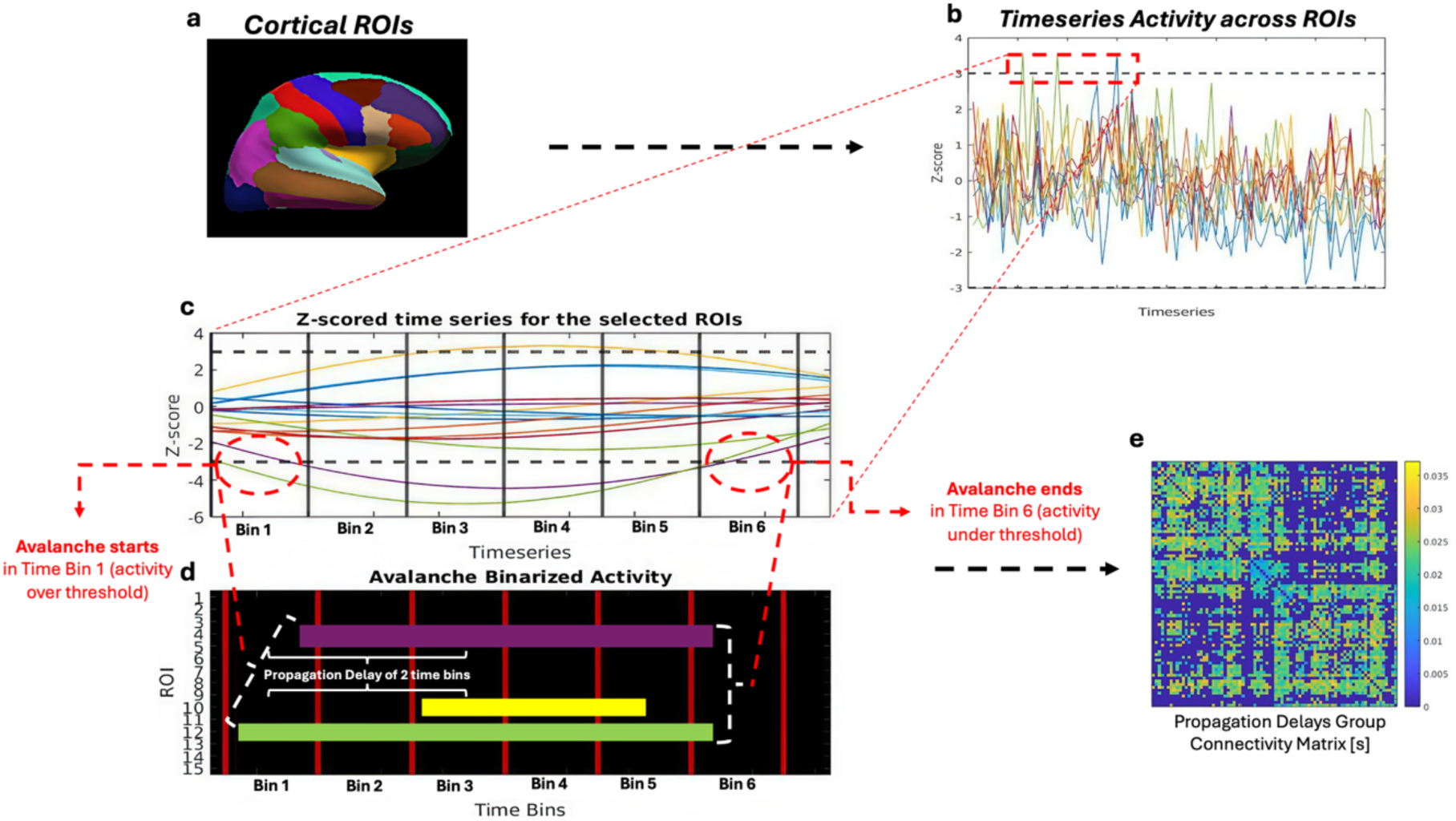
Visual representation of how propagation delays matrices were obtained using the neuronal avalanches framework. Each cortical region of interest (ROI) corresponds to one virtual channel of MEG activity (a). For exemplary purposes in this figure, we have represented only 15 channels (b). Whenever channel activity goes over threshold (black dotted line) such as in the red square (b), an avalanche is initiated. Once there is no overthreshold activity, the avalanche is terminated. For a sample avalanche, the zoomed-in timecourses (c) and the binarised raster plot (d) highlight the ROIs that are involved, from the initial time bin to the final one. To compute the associated delays, the bins between the overall start of the avalanche and any subsequent over-threshold activity are counted for all the ROIs involved. We do this for all avalanches for each subject, and then compute the related average for each pair of ROIs, that are then combined in a group matrix (e).

### Statistical Analysis

For validation purposes, we randomly split our dataset into two using the MATLAB random permutation built-in function – a main dataset of 60 subjects and a test dataset of 34 subjects.

To explore the relationships between the W/GM metrics and the associated delays, we performed Spearman’s correlations (due to some of the variables not being normally distributed) on the main dataset. All eight correlations were corrected for multiple comparisons (8) using the Bonferroni correction (Dunn, 1961). For the WM metrics relationships only, we also conducted a mediation analysis in R to explore whether tract length mediated the associations between each WM metric and propagation delays. For each of the mediation analyses we performed 500 bootstrapping simulations to estimate the indirect effect of the mediator variable and to derive confidence intervals for our data.

To identify which combination of WM and GM variables best predicts propagation delays, we fitted four linear regression models to the main dataset. To ensure that we preserved the expected functional form of the microstructural metrics in relation to the propagation delays, we used the inverse of the axonal diameter, g-ratio and SANDI neurite signal fraction variables. For WM metrics, the expected relationships were derived from Rushton’s proposed model of conduction (Rushton, 1951), while the GM-specific relationship was derived from theoretical considerations: according to cable theory (Rall, 1962), thicker dendrites lead to lower axial resistance and therefore faster propagation. All variables used in the regression models were z-score normalised. To judge the performance of our models we used the square root of the mean squared error 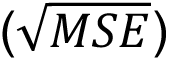 and the adjusted R^2^.

We first modelled tract length alone given its prominent association with delays. We then added WM microstructural metrics alongside tract length to assess their incremental contribution. In a parallel model, we added GM microstructural metrics alongside tract length. Finally, we combined all WM and GM metrics (together with tract length) in a single model. As an external validation, we applied the fitted specifications from these four models to the independent test dataset. Results from the four models in the main set and their counterparts in the test set were adjusted for multiple comparisons using Bonferroni correction (Dunn, 1961).

While for the previously described analyses we explored the MEG whole signal (1-48 Hz) bandwidth delays, we also investigated how the different MEG frequency band delays relate to WM tract length. We performed Spearman’s correlations on the main dataset. All five correlations were corrected for multiple comparisons using the Bonferroni correction (Dunn, 1961).

Finally, to explore the relationships between the different MEG-derived frequency band delays and the WM and GM structural metrics we performed a canonical correlation for the main set of 60 subjects. The W/GM structural variable set consisted of the z-score normalised group level matrices of SF, NF, SR, g-ratio, axonal radius and tract length. The delay matrices consisted of the z-score normalised group-level matrices for the 5 frequency bands (delta, theta, alpha, beta and gamma). We used Wilks’ lambda to assess the significance of each of the relationships between the two sets and we finally used Bartlett’s chi-square approximation to compute the respective p values using a significance level of 0.05. To assess whether the contribution of each of the canonical loadings to the observed relationships was significant we used bootstrapping over 2000 simulations.

## Results

### Whole Signal Propagation Delays and White Matter Relationships (Main Set)

We found a significant positive relationship between tract length and the whole signal delays ρ = 0.44, p<0.0001, implying that longer tracts are associated with longer delays (Figure 3a). Counterintuitively, we also found a significant positive relationship between axonal radius and the delays ρ = 0.44, p<0.0001 (Figure 3b). Interestingly, looking at Figure 3b. we can see that this relationship seems to be mediated by tract length with longer tracts with larger radiuses having longer delays. The mediation analysis revealed a significant indirect effect of tract length, suggesting that WM pathway length mediates the relationship between axonal radius and the observed delays b= 0.01, 95% CI [0.008, 0.013], p<0.001. After accounting for tract length’s mediating effect, the direct effect of axonal radius remained statistically significant b= 0.014, 95% CI [0.010, 0.017], p<0.001. The total effect of axonal radius on the propagation delays was significant b= 0.024, 95% CI [0.021, 0.026], p<0.001. Overall, tract length accounted for 43% of the total observed effect.

**Figure 3.**
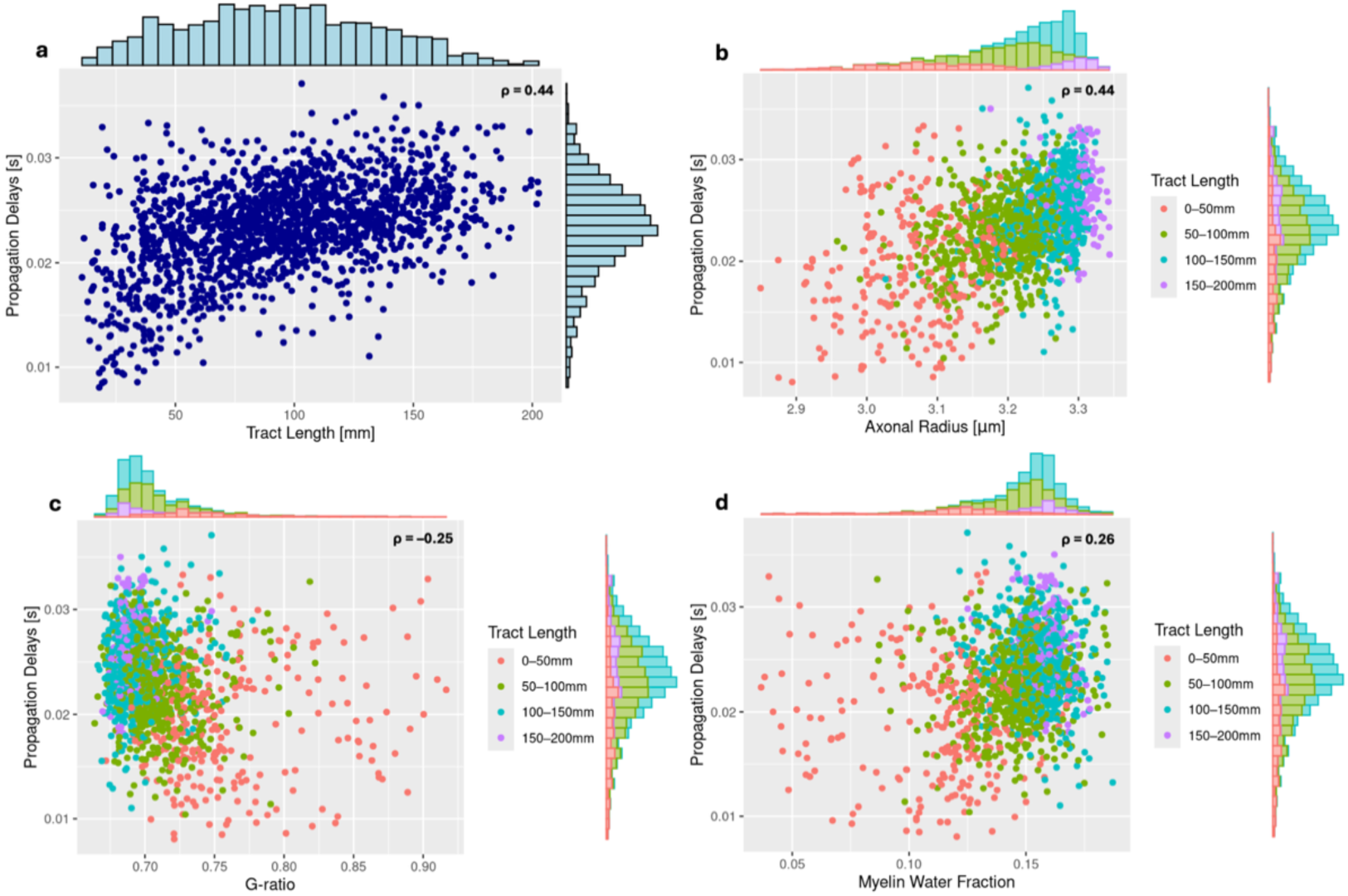
Relationships between WM metrics and the MEG-derived whole signal propagation delays. For each WM variable a histogram of the distribution of values is presented on the corresponding axes. For panels b, c and d the data points are coloured by tract length. For all panels we have plotted only the data points for which the corresponding delays and tract length are nonzero. The top right corner of each panel contains the correlation coefficient (ρ) for each relationship.

When examining the myelin-related metrics, we observed a significant negative association between g-ratio and propagation delays (ρ = −0.25, p < 0.0001; Fig. 3c) and a significant positive association between MWF and delays (ρ = 0.26, p < 0.0001; Fig. 3d). In both cases, the relationships showed a dependence on tract length, similar to that seen for axonal radius. Together, these patterns indicate that connections with shorter tract lengths tend to exhibit lower myelination than longer tracts.

The g-ratio mediation analysis revealed a significant indirect effect of tract length, supporting the idea that length mediates the relationship between g-ratio and the delays b=-0.035, 95% CI [-0.003, 0.013], p<0.001. After accounting for tract length’s mediating effect, the direct effect of g-ratio disappeared b= 0.005, 95% CI [0.010, 0.017], p>0.005. The total effect of g-ratio on the propagation delays was significant b=-0.031, 95% CI [-0.038,-0.022], p<0.001. Tract length fully accounted for the total observed effect, indicating that tract length fully mediated the relationship between the g-ratio estimates and the delays.

The MWF mediation also demonstrated a significant indirect effect of WM length on the observed delays b=0.058, 95% CI [0.051, 0.066], p<0.001. Accounting for the mediating effect of length, the direct effect of MWF on the relationship was not significant b=-0.002, 95% CI [0.016, 0.011], p>0.005. The total effect of MWF was significant b= 0.056, 95% CI [0.043, 0.068], p<0.001. Overall, tract length fully accounted for the total observed effect, implying that tract length fully mediated the relationship between the MWF and the delays.

### Whole Signal Propagation Delays and Grey Matter Relationships (Main Set)

We found significant negative correlations between NF and SR and the delays ρ = - 0.21, p<0.0001 and ρ =-0.18, p<0.0001 respectively (Figure 4a. and 4c.). We also found significant positive relationships between SANDI’s SF ρ = 0.14, p<0.0001, CT ρ = 0.11, p<0.0001 and the corresponding propagation delays (Figure 4b. and 4d.). These results suggest that propagation delays increase with CT and SF, while decreasing with NF and SR. We observed comparable trends for the apparent number of cells and the apparent total surface area (Figure S1).

**Figure 4.**
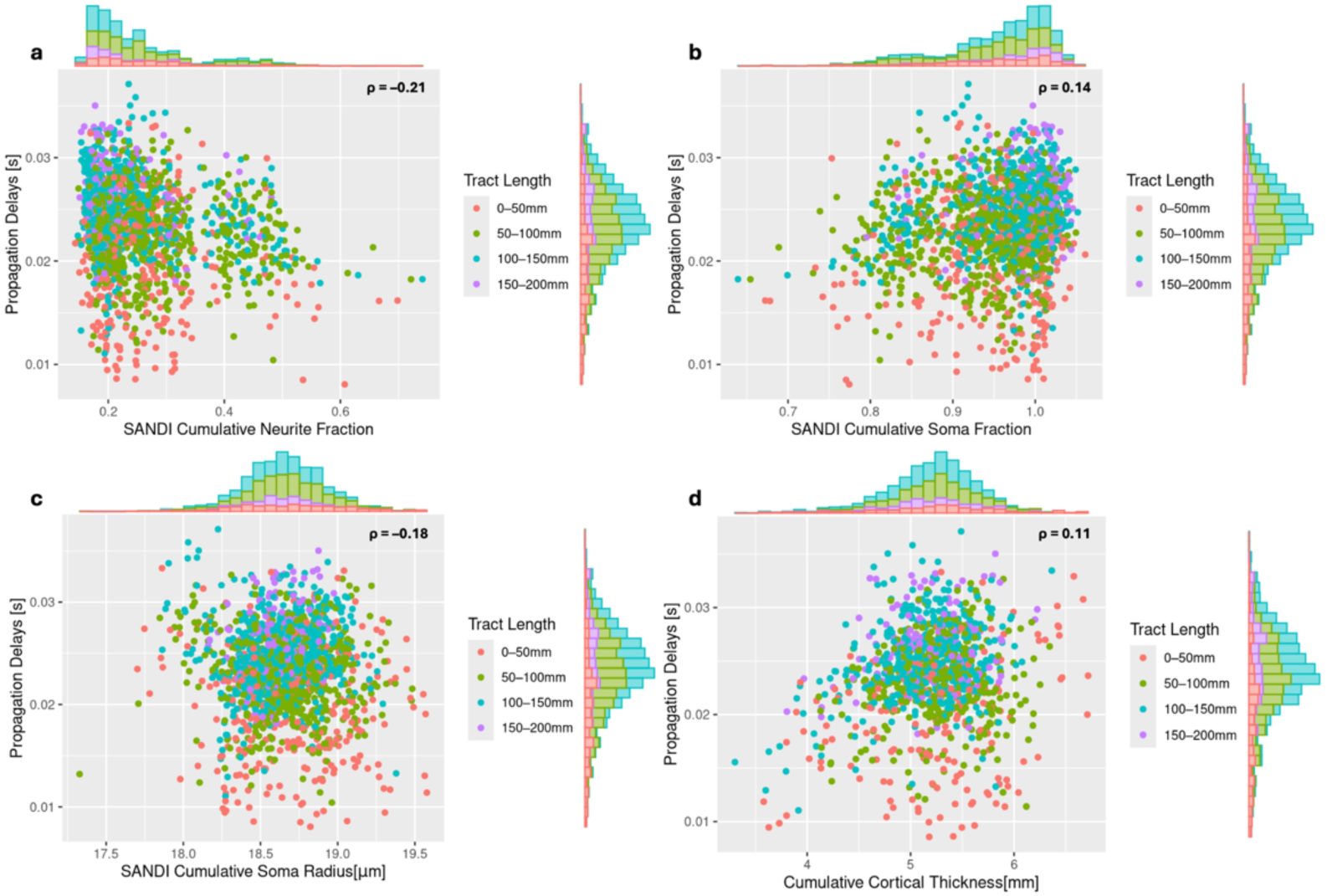
Relationships between GM metrics and the MEG-derived whole signal propagation delays. For each GM variable we summed the average values for two interconnected regions of interest where an avalanche was detected. For each GM metric a histogram of the distribution of values is presented on the corresponding axes. In all panels the data points are coloured by tract length. Panel d includes only cortical regions. For all panels we have plotted only the data points for which the corresponding delays and tract length are nonzero. The top right corner of each panel contains the correlation coefficient (ρ) for each relationship.

### Linear Regression Models

Firstly, we attempted to predict the observed delays using only tract length as a predictor. Table 1. demonstrates that tract length alone accounts for around 22% of the observed delays’ variance. Following this we included the axonal diameter and g-ratio in our model, which improved its’ performance, reflected in the reduction of the square root of the mean squared error (MSE) and increase in explained variance of ~25% (Table 2.).

**Table 1.** Main set Model 1: *Propagation Delays ~ Tract Length*.

| | Estimate | Standard Error | T stat | P value | $\sqrt{MSE}$ | Adjusted R<sup>2</sup> |
| --- | --- | --- | --- | --- | --- | --- |
| **Tract Length** | 0.467 | 0.015 | 30.676 | 1.6379e-182 | 0.884 | 0.218 |

**Table 2.** Main set Model 2: 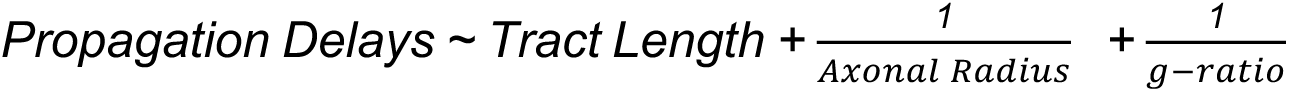.

| | Estimate | Standard Error | T stat | P value | $\sqrt{MSE}$ | Adjusted R<sup>2</sup> |
| --- | --- | --- | --- | --- | --- | --- |
| **Tract Length** | 0.293 | 0.024 | 12.152 | 2.7365e-33 | 0.864 | 0.253 |
| $\frac{1}{\text{Axonal Radius}}$ | -0.312 | 0.025 | -12.538 | 2.7965e-35 | | |
| $\frac{1}{g - ratio}$ | -0.114 | 0.02 | -5.805 | 7.0276e-09 | | |

In parallel we investigated how the GM SANDI metrics and tract length are able to capture the propagation delays (Table 3.). Compared to our WM metrics regression, here we only had two significant contributors, namely tract length and SR, and the regression model performance did not improve on the previous models.

**Table 3.** Main set Model 3: 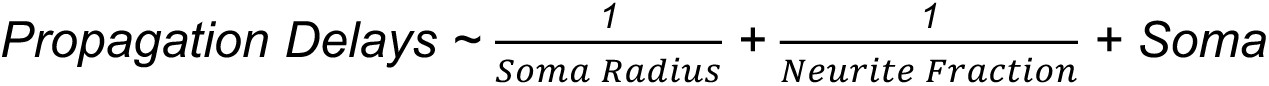 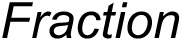.

| | Estimate | Standard Error | T stat | P value | $\sqrt{MSE}$ | Adjusted R<sup>2</sup> |
| --- | --- | --- | --- | --- | --- | --- |
| **Tract Length** | 0.450 | 0.016 | 29.129 | 1.3485e-166 | 0.876 | 0.233 |
| $\frac{1}{\text{Soma Radius}}$ | 0.097 | 0.027 | 3.657 | 2.5919e-4 | | |
| $\frac{1}{\text{Neurite Fraction}}$ | 0.061 | 0.052 | 1.193 | 0.233 | | |
| **Soma Fraction** | -0.036 | 0.040 | -0.897 | 0.370 |  |  |

Finally, we incorporated all the WM and GM variables together in our last regression model (Table 4.). While not all regressors were significant (such as SR), overall, this was the best performing model, although the improvement from the WM regression model was not as substantial.

**Table 4.** Main set Model 4: 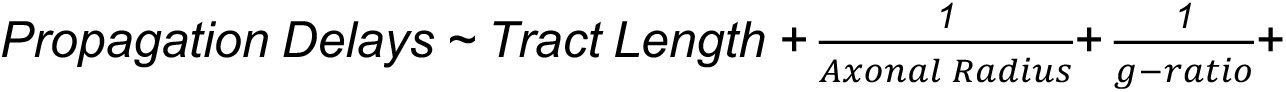 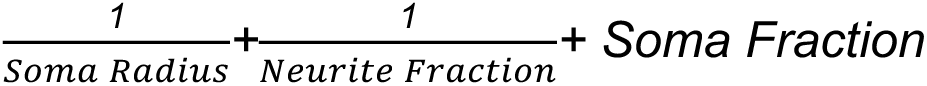.

| | Estimate | Standard Error | T stat | P value | $\sqrt{MSE}$ | Adjusted R<sup>2</sup> |
| --- | --- | --- | --- | --- | --- | --- |
| **Tract Length** | 0.292 | 0.026 | 11.429 | 1.047e-29 | 0.86 | 0.261 |
| $\frac{1}{Axonal Radius}$ | -0.319 | 0.029 | -10.907 | 3.0345e-27 | | |
| $\frac{1}{g - ratio}$ | -0.119 | 0.02 | -6.087 | 1.2796e-09 | | |
| $\frac{1}{Soma Radius}$ | 0.047 | 0.027 | 1.764 | 0.078 | | |
| $\frac{1}{Neurite Fraction}$ | 0.106 | 0.051 | 2.084 | 0.037 | | |
| **Soma Fraction** | -0.146 | 0.041 | -3.563 | 0.0004 |  |  |

**Table 5.** Test set model performance when applying the learned models from our main dataset to our test dataset. Column one demonstrates the used model; column two represents the square root of the mean square error when the main dataset model has been applied to the test dataset. Column three shows the R squared, which demonstrates the proportion of variability, explained by the regressors.

| | $\sqrt{MSE}$ | R<sup>2</sup> |
| --- | --- | --- |
| **Model 1** *Propagation Delays ~ Tract Length* | 1.033 | 0.157 |
| **Model 2** *Propagation Delays ~ Tract Length + $\frac{1}{Axonal Radius}$ + $\frac{1}{g-ratio}$* | 1.018 | 0.181 |
| **Model 3** *Propagation Delays ~ Tract Length + $\frac{1}{Neurite Fraction}$ + Soma Fraction + $\frac{1}{Soma Radius}$* | 1.030 | 0.161 |
| **Model 4** *Propagation Delays ~ Tract Length + $\frac{1}{Neurite Fraction}$ + Soma Fraction + $\frac{1}{Soma Radius}$ + $\frac{1}{Axonal Radius}$ + $\frac{1}{g-ratio}$* | 1.016 | 0.184 |

Following this, we applied regression models fitted on the main set to our test dataset for validation. The test set demonstrated explained variance and mean error values in line with the main dataset, suggesting that the models are generalising.

### MEG Frequency Bands Propagation Delays’ Association with Tract Length

Starting with the slowest frequency band, we found a significant negative correlation between delta band propagation delays and tract length ρ =-0.15, p<0.0001 (Figure 5a.). For the theta band delays and tract length we found a significant positive relationship ρ = 0.18, p<0.0001 (Figure 5b.). For alpha and beta frequency bands we also found significant positive correlations with length ρ = 0.27, p<0.0001 and ρ = 0.40, p<0.0001 respectively (Figure 5c. and 5d.). Finally, for the gamma band we found a significant negative relationship with tract length ρ =-0.25, p<0.0001 (Figure 5e.). Overall, it seems that there is a trend in the length-delay relationships - from slower to faster frequency bands the relationship seems to become stronger in absolute value. This trend, however, is not apparent in the delta and gamma bands. To aid the interpretation of our results the mean number of avalanches for each frequency band is reported in Table S1.

**Figure 5.**
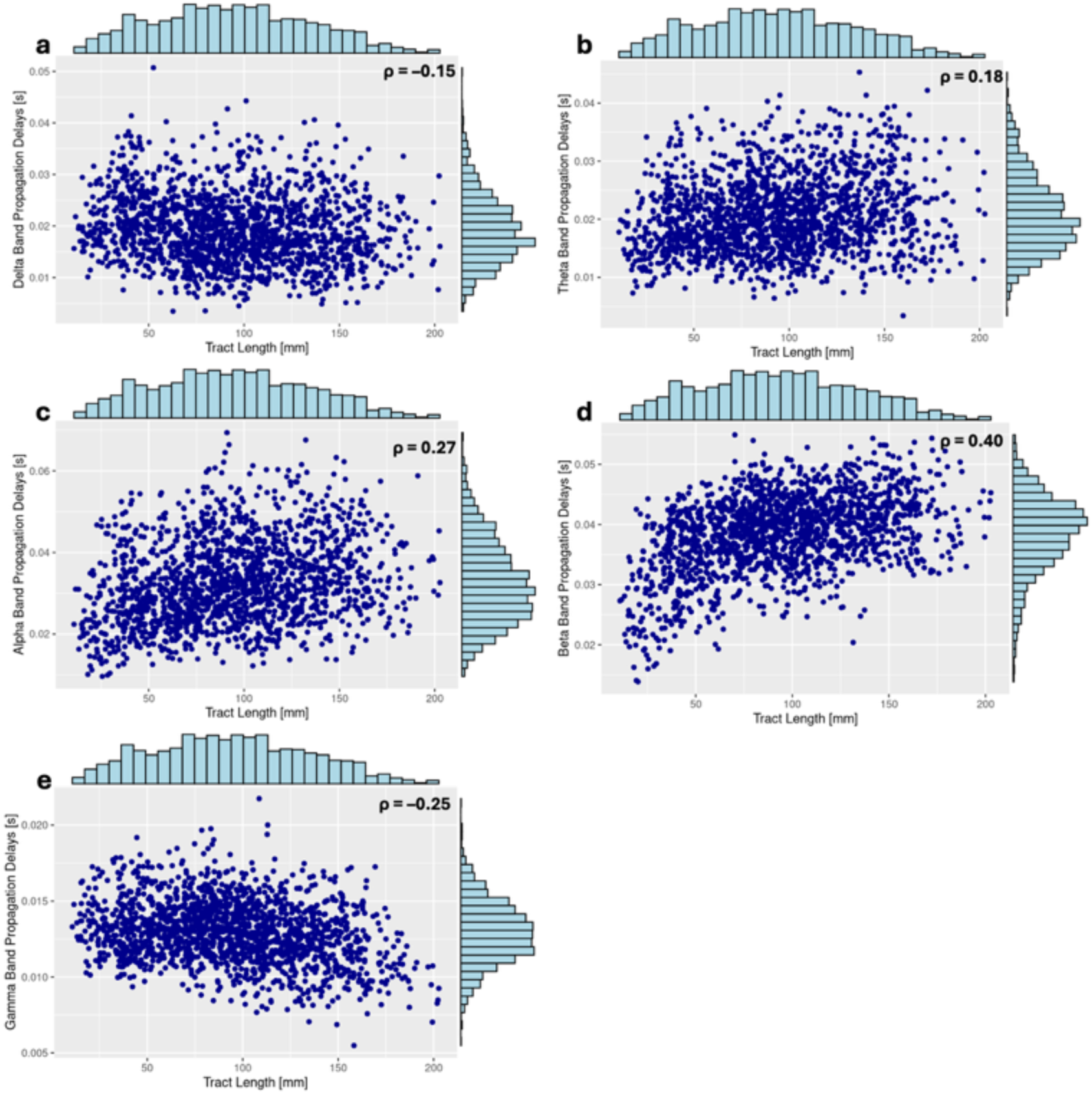
Relationships between WM tract length and the different MEG-derived frequency bands delays in each of the different panels. For each frequency band a histogram of the distribution of values is presented on the corresponding axes. For all panels we have plotted only the data points for which the corresponding delays and tract length are nonzero. The top right corner of each panel contains the correlation coefficient (ρ) for each relationship.

### Canonical Correlation Analysis (CCA)

The overall CCA model revealed three significant CCA pairs. Figure 6. shows the canonical loadings for these 3 pairs and their respective r and p values. For the first canonical variate pair, the structural variables contributing most strongly were tract length (loading = 0.966), 95% CI [0.779, 0.993], SF (loading= 0.690), 95% CI=[0.493, 0.825], axonal radius (loading=0.408), 95% CI=[0.154, 0.590] and g-ratio (loading=0.586), 95% CI=[0.279, 0.788], while the dominant contributors on the delay side were alpha (loading=0.545), 95% CI=[0.222, 0.754], beta (loading=0.924), 95% CI=[0.716, 0.986], delta (loading=0.559), 95% CI=[0.214, 0.781] and gamma (loading=0.902), 96% CI=[+0.616, +0.987]. This canonical pair reflects a relationship whereby increased microstructural metrics values are associated with increased propagation delays across multiple frequency bands. For the second and third canonical pairs, there were no individual microstructural or delay variables that were significant contributors to the observed relationships. Therefore, in these two roots there were no robust relationships between the two sets of variables.

**Figure 6.**
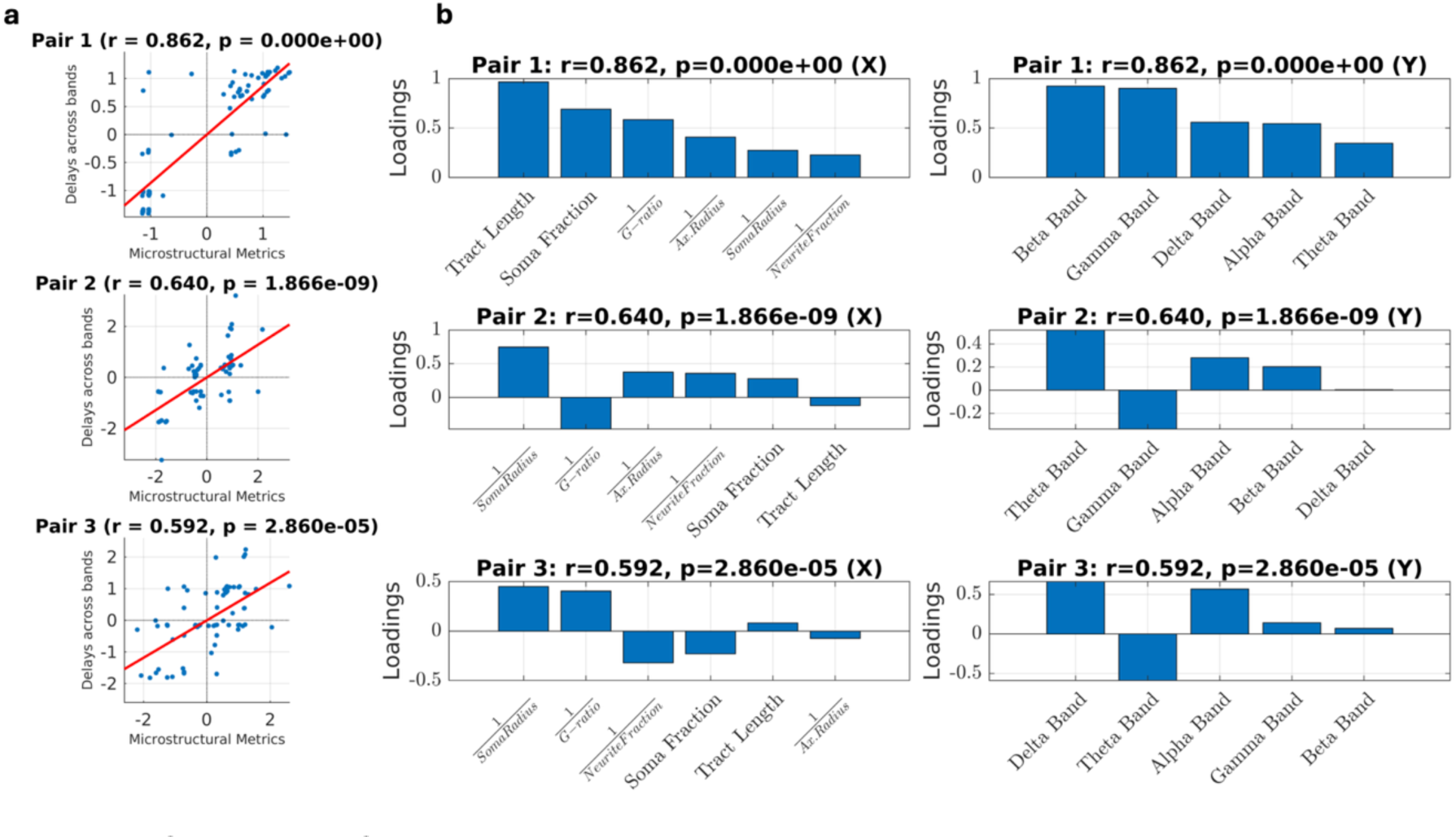
Canonical Correlation Analysis. Panel a shows the canonical correlations between the MRI metrics and the MEG delays for each of the significant pairs. Panel b represents the canonical loadings and the respective r- and p-values for each significant pair. The left side represents the microstructural variables, while the right side represents the different frequency band delays and their canonical loadings.

## Discussion

To gain a better understanding of how structure is associated with large-scale neural dynamics, we explored the relationships between MEG-derived propagation delays and a range of MRI WM and GM structural metrics. Building upon this, we also explored how well different combinations of WM and GM metrics predicted whole signal propagation delays. We then investigated frequency-specific propagation dynamics by examining the relationships between delays across the five different MEG functional frequency bands and tract length. Finally, using canonical correlation analysis, we demonstrated multivariate coupling between WM and GM structural metrics and MEG frequency band-specific propagation delays.

Consistent with prior work on WM specific conduction delays, we observed a positive relationship between pathway length and propagation delays, suggesting that longer WM connections are associated with increased propagation times (Caminiti et al., 2009; Tomasi et al., 2012).

In myelinated axons, conduction velocity is thought to be directly proportional to axonal diameter, implying that larger axons support faster propagation and therefore shorter delays (Waxman, 1980). At first glance, our finding of a positive relationship between axonal radius and whole-signal propagation delays seems counterintuitive. Similarly, although higher myelination is classically associated with faster conduction and shorter delays, our myelin metrics show the opposite trend (Waxman, 1980). This discrepancy may be explained by the strong dependence of axonal radius, g-ratio and MWF on tract length. Our results (also confirmed by the mediation analysis) suggest that longer WM pathways contain tracts with larger axonal radius and higher myelination, potentially reflecting the need for optimized structure to support timely long-range conduction. Therefore, although larger individual myelinated axons may generally support faster conduction, longer anatomical pathways at the bundle level could lead to a positive relationship between axonal radius, MWF and propagation delays at the macroscale. Furthermore, since we used COMMIT to prune implausible streamlines, the resulting tract-specific axonal-radius estimates likely favour anatomically plausible long connections. This may accentuate the observed coupling between larger axonal radius and longer pathways at the macroscale.

To our knowledge the relationship between SANDI metrics and propagation delays has not been explored previously. As mentioned, there have been physiological attempts such as Rall’s cable theory to model dendritic properties and how they shape how input signals are propagated and integrated throughout a neuron (Rall, 1962). Similarly to the axons, the morphological properties of dendrites such as their branching, diameter and voltage-gated ion channels’ activity have been implicated in actively shaping signal transmission dynamics (Stingl et al., 2025). Cable theory treats dendrites as electrical cables and explicitly predicts that larger dendritic diameters reduce axial resistance, thereby facilitating faster signal propagation. Such morphological properties would likely be reflected primarily by NF, suggesting that higher NF values should be associated with shorter propagation delays, consistent with our findings.

The size and morphology of neuronal somata influence key electrical properties of neurons, including membrane resistance, capacitance, and integrative time constants, with larger somas generally exhibiting lower input resistance and reduced excitability (Henneman et al., 1965; Dukkipati et al., 2018). From a cable-theory perspective, neurons can be approximated as distributed electrical systems in which membrane and axial resistances jointly determine the efficiency and timing of signal propagation across cellular compartments (Rall, 1962). In this framework, somatic surface area contributes to membrane capacitance and current leakage, thereby shaping the temporal dynamics of voltage integration. Within the SANDI model, these properties are indirectly reflected in FS and SR, which together describe the contribution of restricted spherical compartments to the diffusion signal. However, SF does not uniquely index soma size, as it is jointly influenced by soma density and morphology within a voxel, and may therefore reflect different underlying microstructural configurations (e.g., few large somas versus many smaller somatic compartments). This inherent ambiguity in SF may partly explain the difficulty in establishing a clear and monotonic relationship between soma-related SANDI metrics and propagation delays, despite theoretical expectations regarding the role of somatic geometry in shaping neuronal time constants. Using measures that specifically reflected apparent total surface area and number of cells did not lead to clearer relationships (Figure S1). While these parallels between the SANDI metrics and physiological models may be an oversimplification, they offer a useful first step toward reconciling the GM physiological models with the GM microstructural models.

Finally, cortical thickness is a feature reflective of all cells’ arrangement (He et al., 2007). The observed weak positive relationship might be explained by the heterogeneous nature of cortical thickness, which renders its relationship to propagation difficult to interpret.

In our attempts to predict propagation delays we were able to explain up to ~26% of the observed delays. Our results suggest that propagation delays are associated with both large-scale anatomy and local microstructure, with WM tract length capturing the geometric distance signals must travel, and the remaining metrics reflecting regional tissue and cellular architecture that may modulate effective transmission and integration times. It is important to note that propagation delays are influenced by multiple factors, some of which are discussed below, that are not accounted for in the present analysis.

The neuronal avalanches framework captures large-scale criticality dynamics and can provide insight into how the structural connectome is associated with brain dynamics across temporal scales (Sorrentino et al., 2021). The relationship between MEG frequency band delays and WM tracts reflects how anatomical pathways’ distance shapes differently temporal dynamics. To further explore the relationships between MEG frequency band propagation delays and the microstructural metrics, we performed a CCA. The CCA revealed coordinated structure-function relationships across multiple spatial and temporal scales, suggesting multivariate coupling between WM and GM structural metrics and delays. Together, these findings suggest that both macroscopic anatomical architecture and microscopic tissue properties contribute to shaping the temporal organization of large-scale brain dynamics.

With diffusion MRI we are mostly sensitive to the largest axons within a voxel (Veraart et al., 2020). Histological studies have reported that in human WM most of the axonal diameter values range between 0.5−2 µm, with a very small fraction having a larger diameter than 3 µm (Aboitiz et al., 1992; Caminiti et al., 2009; Veraart et al., 2020). Similarly, we are limited to the tail of the distribution when obtaining the SANDI cell body radii as larger cell bodies have a larger contribution to the diffusion signal. Although the use of an ultra-strong gradient Connectom system increases sensitivity to microstructural features such as smaller axons and somas compared to standard clinical MRI scanners with lower gradient amplitudes, we are still limited by the diffusion resolution limit (Nilsson et al., 2017).

Previous work based on single-axon conduction delay simulations has been able to explain around ~85% of conduction velocity variability using only WM microstructural parameters that could be estimated with MRI (Drakesmith et al., 2019). Beyond these metrics, there were other parameters which are determinants of conduction velocity and delays such as the nodes of Ranvier and their spatial spacing, the electrical properties of the membranes of the axons and myelin sheath, all integral in supporting timely conduction but still currently an unresolved challenge in non-invasive imaging (Waxman, 1980; Seidl, 2014; Drakesmith et al., 2019). Similarly, with the GM metrics - we might be missing crucial information such as the ion channels’ ratio, the dendritic branching structure etc. which have also been implicated in affecting propagation (Stingl et al., 2025). Even with advanced imaging techniques and ultra-strong gradients we are still limited in our ability to measure all the relevant microstructural metrics to model delays.

Since this was the first attempt to relate structural metrics to macroscale delays, we implemented linear regression as the most intuitive statistical approach, given the simplicity of the model and its minimal number of assumptions. We accounted for the functional form of our variables, based on the predictions of the implemented biophysical models in relation to predicting propagation delays e.g., we used 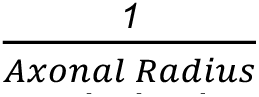 to predict the delays as described in Rushton (1951). While alternative statistical methods might be worth exploring in the future, further assumptions and related supporting evidence may be needed.

For the purposes of this work, we summed the GM indices of connected GM ROIs where an avalanche was detected. However, this approach lacks biological specificity, as the two connected regions are likely to differ their morphology, organisation and properties (Zeng & Sanes, 2017) and their contributions to delays may not just simply cumulate. Future work should characterise GM microstructure more precisely, for example by further disentangling avalanche structure and examining avalanches during task rather than resting state. This would allow the study of more isolated brain systems, such as the motor system, which are better described histologically and would therefore increase the interpretability of our results.

In conclusion, our findings suggest that both large-scale structure and local tissue microstructure are associated with large-scale dynamics, providing a foundation for future work non-invasively investigating the biological underpinnings of large-scale neural transmission.

## Acknowledgements

DKJ and the WAND dataset acquisition were supported by a Wellcome Trust Investigator Award (096646/Z/11/Z) and a Wellcome Trust Strategic Award (104943/Z/14/Z). M.P. was supported by the UKRI Future Leaders Fellowship (MR/T020296/2). MM was funded by the Wellcome Trust through a Sir Henry Wellcome Postdoctoral Fellowship (213722/Z/18/Z).

