## Supplementary Materials Figure S1. and Table S1. for "From Brain Microstructure to Dynamics: Linking Grey and White Matter Architecture to Propagation Delays"

Estimating the apparent number of cells and the total surface from SANDI

The SANDI model can be used to compute alternative measures of the underlying cell populations (Carriero et al., 2026), in particular the apparent number of cells and the apparent total surface area, respectively calculated as:

$$f_{c}=\frac{Vf_{s}}{\frac{4}{3}\pi r_{s}^{3}}$$

$$f_{sup}={Vf}_{c}\cdot4\pi r_{s}^{2}$$

where V is the voxel volume. For each ROI, we computed f_c_ and f_sup_ and then, in line with what was done with the other GM metrics, for each pair of ROIs we calculated the respective cumulative value, building the related connectivity matrices.

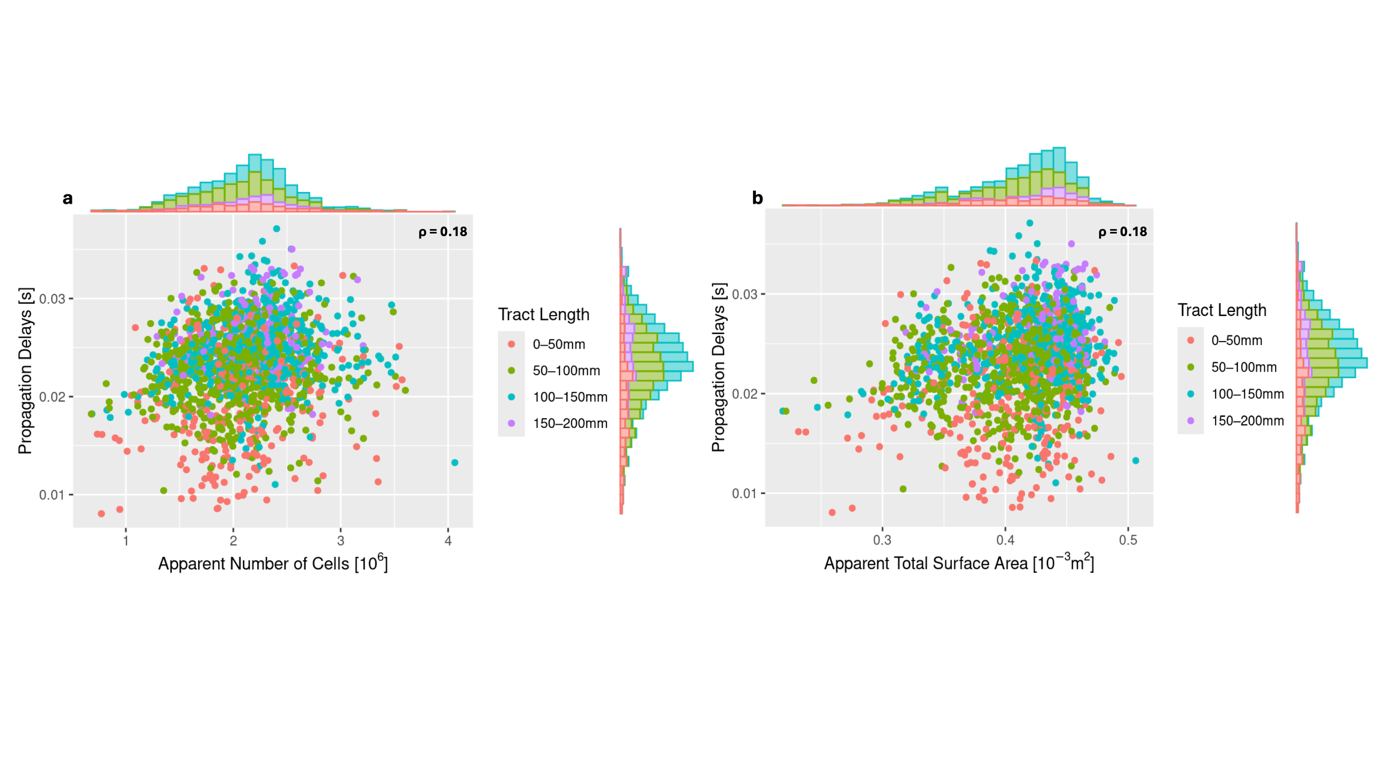

**Figure S1**. Relationships of the cumulative SANDI-based apparent number of cells and total surface area with MEG-derived whole signal propagation delays, highlighting through histograms the distribution of each metric and using color coding to represent different tract lengths.

| **Frequency Band** | **Average Number of Avalanches** | **Standard Deviation** |
| --- | --- | --- |
| Whole Signal (0.5-48Hz) | 2042.1 | 336.389 |
| Delta (1-4Hz) | 1253.5 | 118.566 |
| Theta (4-8Hz) | 1969.3 | 216.895 |
| Alpha(8-13Hz) | 990.417 | 138.002 |
| Beta(13-30Hz) | 1474.2 | 119.712 |
| Gamma(30-48Hz) | 769.083 | 137.190 |

**Table S1**. The average number (and standard deviation) of detected avalanches for each tested frequency band for the main set of 60 subjects.
